# Multiplexity and Graph Signal Processing of EEG Dynamic Functional Connectivity Networks As Connectomic Biomarkers for Schizophrenia Patients: A Whole Brain Breakdown

**DOI:** 10.1101/551671

**Authors:** Stavros I. Dimitriadis

**Affiliations:** Cardiff University Brain Research Imaging Centre, School of Psychology, Cardiff University, Cardiff, United Kingdom; Neuroinformatics Group, Cardiff University Brain Research Imaging Centre, School of Psychology, Cardiff University, Cardiff, United Kingdom; Division of Psychological Medicine and Clinical Neurosciences, School of Medicine, Cardiff University, Cardiff, United Kingdom; School of Psychology, Cardiff University, Cardiff, United Kingdom; Neuroscience and Mental Health Research Institute, School of Medicine, Cardiff University, Cardiff, United Kingdom; MRC Centre for Neuropsychiatric Genetics and Genomics, School of Medicine, Cardiff University, Cardiff, United Kingdom

**Keywords:** EEG, dynamic functional connectivity graphs, multiplexity, chronnectomics, dominant intrinsic coupling modes, schizophrenia

## Abstract

**Introduction:** Last years, many studies explored the disruption of functional interactions in schizophrenic patients compared to healthy controls supporting the consideration of schizophrenia as a disconnection syndrome. However, the majority of studies followed a static connectivity analysis ignoring the rich information encapsulated under the framework of dynamic functional connectivity graph (dFCG) analysis.

**Methods:** A dFCG has been estimated using a multivariate phase coupling estimator (PCE) by integrating both intra and cross-frequency coupling modes into an integrated DFCG (iDFCG) that encapsulates the functional strength and the type of coupling between pairs of brain areas. We analysed dFCG (Low-Order iDFCG) profiles of electroencephalographic resting state (eyes closed) recordings of healthy controls (n=39) and subjects with symptoms of schizophrenia (n=45) in basic frequency bands {δ,θ,α_1_,α_2_,β_1_,β_2_,γ}. We constructed the High-Order - iDFCG by adopting the cosine similarity between the time-series derived from the Low-Order-iDFCG. Estimating Graph Laplacians transformations of Low-Order and High-Order-iDFCG and by calculating the temporal evolution of Synchronizability (Syn), network metric time series (NMTS^Syn^) were produced.

**Results:** Following, a machine learning approach based on multi-kernel SVM with the four NMTS^Syn^ used as potential features and appropriate kernels, we succeeded a superior classification accuracy (∼98%). DICM and flexibility index (FI) achieved a classification with absolute performance (100 %).

**Conclusions:** Schizophrenic subjects demonstrated a hypo-synchronization compared to healthy control group which can be interpreted as a low global synchronization of co-fluctuate functional patterns. Our analysis could be helpful for clinical prediction of ScZ and also for evaluating non-intervention treatments tailored to schizophrenia.

## 1. Introduction

Schizophrenia (ScZ) is a mental disorder that is characterized by strange speece, abnormal behaviour and decreased ability to understand reality (American Psychiatric Association,2013). Last decade, neuroimaging and network neuroscience explores how ScZ alters brain activity (Hunt et al., 2017) and connectivity (Kambeitz et al., 2016). These findings defined ScZ as a brain disconnectivity disorder (Schmitt et al, 2011; Friston,1996) characterized by an aberrant fronto-parietal interactions compared to healthy controls (Friston and Frith, 1995). Animal models proved that lesions in the hippocampus may cause a disconnected functional connectivity pattern in the prefrontal cortex (Schmitt et al, 2011). Magnetic resonance imaging (MRI) studies tailored to schizophrenic patients revealed deficits of gray matter in medial temporal lobe (and hippocampus), in the prefrontal, anterior cingulate, parietal and superior temporal cortex and the thalamus and a magnification of the ventricles (Shenton et al., 2001). Diffusion tensor imaging (DTI) studies have demonstrated a decrement of white matter anisotropy in the temporal and left prefrontal cortex (Kubicki et al., 2007). It is important to underline current evidences that limbic pathways connecting thalamus, anterior cingulate cortex, prefrontal cortex and the hippocampus are activated in information processing and higher cognition (Gaebel,2011). Higher cognition refers to cognitive processes that presuppose the availability of knowledge that put it in use. A first clinical picture of the aberrant connectivity in schizophrenia may be the decreased of functional integration resulting in the malfunction of association process (Gaspar et al., 2008).

Cortical disconnectivity theory for schizophrenia has been explored also with functional neuroimaging. The analysis of multichannel electro (EEG) and magneto-encephalography (MEG) signals provides unique information for functional interaction between neural oscillations characterizing brain activity of specific areas within the same frequency (intra-frequency coupling) and between frequencies (cross-frequency coupling; Dimitriadis et al., 2017b,2018c) in high temporal resolution (Canolty and Knight, 2010; von Stein and Sarnthein, 2000; Lisman and Buzsaki, 2008; Dimitriadis et al., 2017b, 2018c). These functional interactions have been proved to be altered in schizophrenia (Allen et al., 2011). The analysis of EEG at resting state condition demonstrated and increased pattern in δ (1 – 4 Hz), θ (4 – 8 Hz) and β (13 – 30 Hz) frequencies complementary to a decreased pattern in α frequency within the frontal cortex (“*hypo-frontality*”) in schizophrenic patients relative to controls (Ingwar and Franzen, 1974; Ragland et al., 2007). An additional potential biomarker for schizophrenia is *abnormal brain asymmetry* (Oertel-Knoechel et al., 2012; Gotts et al., 2013; Ribolsi et al., 2014; Miyata et al., 2012).

A recent EEG study at resting-state explored aberrant static functional brain connectivity induced by schizophrenia adopting three connectivity estimators, alternative network metrics and two reference systems (Olejarczyk and Jernajczyk, 2017). The whole connectivity analysis focused on five frequencies: 2–4 Hz (δ), 4.5–7.5 Hz (θ), 8–12.5 Hz (α), 13–30 Hz (β), 30–45 Hz (γ). They revealed the inter-hemispheric asymmetric group-difference using Directed Transfer Function granger causality estimator at resting-state. Another study proposed a methodology of selecting the best set of EEG sensors based on their connectivity profile with the rest of EEG sensors in order to design an optimal classifier for the discrimination of healthy controls (thirst) from schizophrenic patients (thirty-two) (Dvey-Aharon et al., 2017). Connectivity analysis has been realized with correlation between pair-wise time series in a broadband frequency of 0.1 Hz-30 Hz. They finally reported a classification performance of 93.8%. The main disadvantage of both studies was the static connectivity analysis approach versus a more dynamic that takes the advantage of the temporal resolution of EEG modality. Moreover, only the first study explored static connectivity analysis within specific frequencies while both didn’t explore also cross-frequency interactions (Dimitriadis et al., 2015a, 2016a,b, 2018a,c,d).

The majority of studies that explored functional connectivity at α frequency band reported a decreased pattern in ScZ. In particular, decreased α connectivity estimated by coherence, lagged coherence and phase synchrony, has been reported at frontal (Di Lorenzo, et al., 2015; Tauscher et al.,1998), fronto-posterior (Di Lorenzo, et al., 2015, Lehmann et al., 2014) and parieto-temporal (Di Lorenzo, et al., 2015) brain areas (for different results see also Andreou et al., 2015, Kam et al., 2013, Winterer et al., 2011, Merrin and Floyd, 1996). Interestingly, two studies reported a high correlation between functional connectivity at rest in α frequency band with symptoms in ScZ (Hinkley et al., 2011; Merrin and Floyd, 1996). Evidences that contradict have been reported in fast oscillations at β (13– 30 Hz) and γ (30–200 Hz) frequencies at rest, including both elevated (Di Lorenzo, et al., 2015), reduced (Kam et al., 2013) and intact (Lehmann et al., 2014, Andreou et al., 2015, Tauscher et al.,1998, Winterer et al., 2011) β frequency band connectivity. Preliminary evidence suggests that β based functional connectivity is influenced by illness progression and clinical symptomatology (Di Lorenzo, et al., 2015). For a systematic review tailored to functional connectivity evidences using EEG in schizophrenia see Maran et al. (2016).

The EEG studies related to ScZ patients so far analysed functional connectivity under the framework of static functional connectivity graphs (FCG) which is constructed by adopting a bivariate or multivariate connectivity estimator that quantifies the signal temporal synchronization across every pair of brain areas. This type of FCG is called low-order (LO) by definition because they characterize EEG signal synchronizations and not high-level inter-regional interactions. A recent study proposed the notion of high-order (HO) FCG by measuring the inter-regional interactions via the similarity between two brain areas’ functional topographic profile (Zhang et al., 2017). They demonstrated that HO-FCG complements to LO-FCG in the preliminary design of a novel biomarker tailored to mild cognitive impairment based on resting-state fMRI activity (Zhang et al., 2017).

We hypothesized that the construction of an integrated DFCG (iDFCG) with the combination of domain intrinsic coupling modes (DICM) which can be either intra or inter-frequency coupling estimations (cross-frequency couplings) will reveal significant features related to ScZ (Dimitriadis et al., 2017b,2018c). We expected that the probability distribution of dominant coupling modes related to δ frequency will be significant higher for ScZ patients (Sharp and Hendren 2007). We assumed that ScZ patients will demonstrate a less flexible global brain pattern compared to healthy controls while their synchronizability will be lost across the whole brain.

Our study is a combination of advanced signal processing and network neuroscience under the framework of time-varying functional connectivity that describes how the relationship between brain areas change across experimental time. This approach has been described as chronnectome from the combination of the words chronos (Hellenic word for time) and connectome that is the mapping of functional interactions between EEG time series into a brain network (graph). The current study explores how the dominant intrinsic coupling modes which are either intra or cross-frequency couplings altered in ScH parients. We also study how chronnectomic features derived from the whole brain synchronizability change in ScH patients. Especially, we adopted a graph fourier transform of LO and HO-iDFCG via a graph laplacian transformation following a previous analytic pathway (Dimitriadis et al., 2018b; Huang et al., 2018).

In Materials and Methods section, we described the data acquisition and details of the EEG dataset, the pre-processing steps including the denoizing with independent component analysis (ICA). The Results section is devoted to describe the results including classification results and significant changes between groups. Finally, the Discussion section includes the discussion of the current research results with future extensions.

## 2. Material and Methods

The subjects were adolescents who had been screened by psychiatrist and divided into a healthy group (*n* = 39) and a group with symptoms of schizophrenia (*n* = 45). Both groups included only school boys. The age of the schizophrenia group ranged from 10 years and 8 months to 14 years. The healthy control group included 39 healthy schoolboys aged from 11 years to 13 years and 9 months. The mean age in both groups was 12 years and 3 months. EEG activity was recorded from 16 EEG channels at resting-state with eyes closed. The electrode positions are demonstrated in Figure 1. The sampling rate is 128 Hz and the recording time was 1 min giving a total of [60 secs x 128] = 7,680 samples. You can download the EEG recordings from the website: http://brain.bio.msu.ru/eeg_schizophrenia.htm. EEG time series were averaged references before preprocessing steps.

**Fig. 1.**
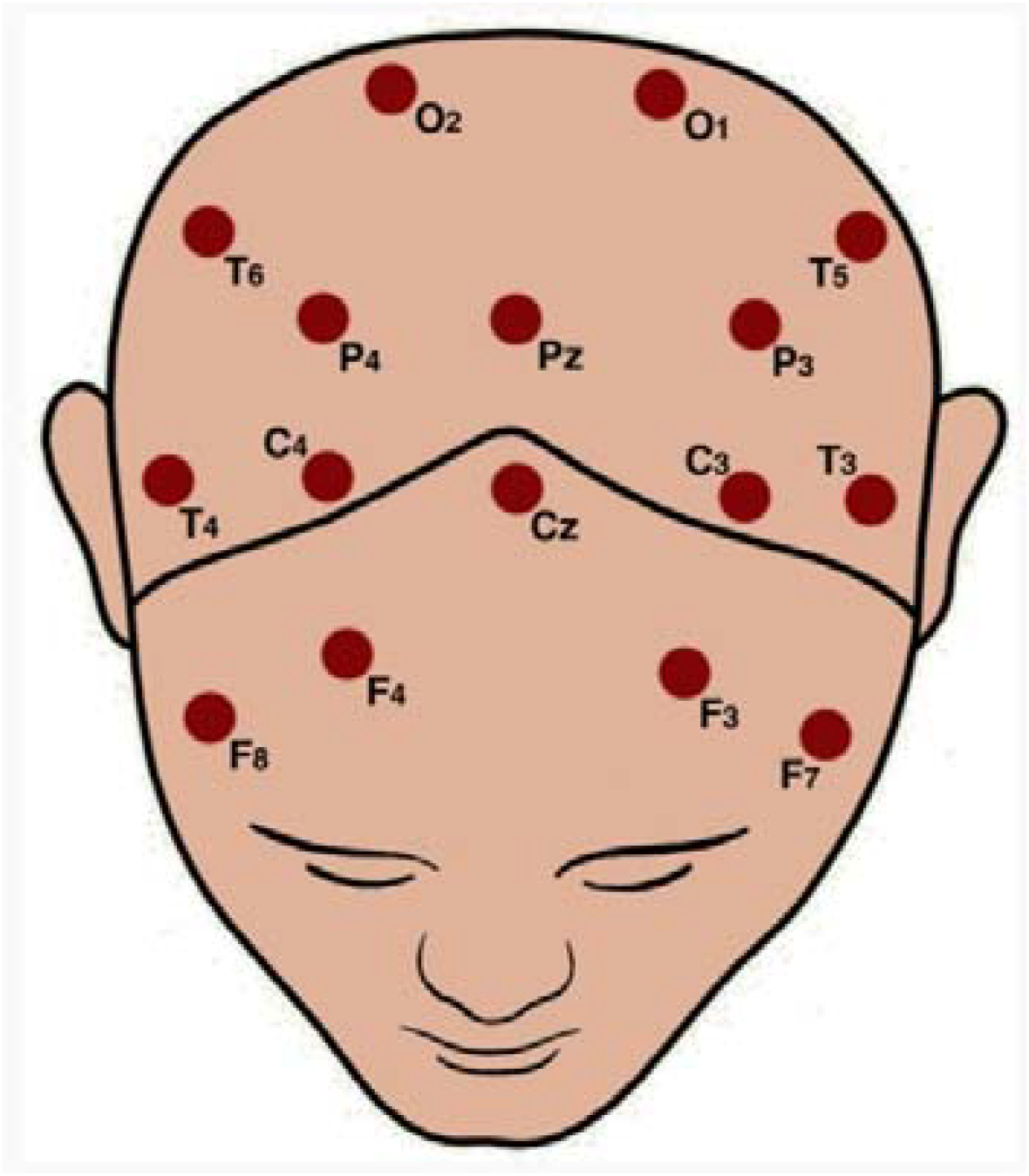
Topology of EEG recording sensors.

A written informed consent was obtained from the parents of the participants in this study.

### 2.1 Artifact Reduction with Independent Component Analysis (ICA)

We cleaned ongoing activity with ICA following preprocessing steps adopted in previous studies of our group (Dimitriadis et al., 2013a,b,2015a,b). We employed the extended Infomax algorithm implemented in EEGLAB (Delorme and Makeig, 2004; see section 1 in supp.material).

### 2.2 Dynamic PCE estimates: the dynamic integrated functional connectivity graph based on PCE (DIFCG^PCE^ graph)

Here, we describe our dominant intrinsic coupling model (DICM) presented in the majority of functional neuroimaging modalities (Dimitriadis,2018c). The goal of the DICM model is to extract the dominant coupling mode between every pair of EEG sensors here and across temporal segments. DICM model defines the dominant coupling mode across intra-frequency phase-to-phase coupling and inter-frequency phase-to-amplitude coupling modes.

We studied dynamic functional connectivity across experimental time and the 16 EEG sensors within and between the seven studying frequency bands {δ, θ, α_1_, α_2_, β_1_, β_2_, γ} defined, respectively, within the ranges {0.5–4 Hz; 4–8 Hz; 8–10 Hz; 10–13 Hz; 13–20 Hz; 20–30 Hz; 30–48 Hz}. EEG recordings were bandpass filtered with a 3rd order zero□phase Butterworth filter using *filtfilt.m* MATLAB function.

The width of the temporal window moved forward across experimental time was set equal to 250ms (or 32 samples), as an adequate number to capture the dynamics of every brain rhythm (fast and slows, Dimitriadis et al.,2013a,b,2015a,b,2016a,b,c,2017a,b,2018b,c). The centre of the temporal window moved forward every 50 ms (6 samples) and both intra and inter□frequency interactions between every all frequencies were re□estimated leading to a ‘quasi□stable’ in time static PCE-based FCG. The multivariate phase coupling estimation (PCE) provides a new approach to reveal the functional coupling based on multivariate phase statistics between nodes in a large network (Cadieu and Koepsell, 2010; Pusil et al., 2019). Here, we repeated the whole analysis by adopting for the very first time a multivariate phase-based estimator instead of a bivariate like the iPLV (Dimitriadis et al., 2018c). This approach leads to 1187 temporal segments.

In this manner, a series of PCE-based graph estimates were computed per subject, for each of the 7 possible intra□frequency coupling refers to the studying frequencies and the 21 possible cross□frequency pairs. This procedure which is described in detail in our previous papers (Dimitriadis and Salis, 2017b; Dimitriadis et al., 2018c), resulted in 7 DFCG^PCE^ per participant for within frequency bands and 21 DFCG^PCE^ per participant for each possible cross□frequency pair. DFCG^PCE^ tabulates PCE estimates between every possible pair of sensors. For each subject, a 4D tensor [frequencies bands (28 × temporal segments × sensors x sources] was created for each subject integrating spatiotemporal phase-based interactions. Afterward, we applied surrogate analysis in order to reveal the dominant type of interaction for each pair of sensors and at each snapshot of the DFCG (see section 2 in sup.material).

Figure 2a illustrates how the DICM is defined in the first two temporal segments from the EEG dataset between F7 and F3 EEG sensors. Finally, we tabulated both the strength and the type of dominant coupling mode in 2 3D tensor [1196 x 16 x 16], one that keeps the strength of the coupling based on PCE and a second one for keeping the dominant coupling of interaction using integers from 1 up to 28:1 for δ, 2 for θ, …, 7 for γ, 8 for δ − θ, …, 28 for β_2_ − γ}. The notion of phase□to□amplitude cross□frequency coupling (CFC) estimator has been employed also in our previous studies with EEG/MEG brain datasets (Dimitriadis and Salis, 2017b, Dimitriadis et al., 2017c,2018a,b,c,d). Figure 2b demonstrates the temporal evolution of DICM across experimental time for F7-F3 EEG pair while the comodulogram on the right side tabulates the PD of the DICM for this semantic time series.

**Figure 2.**
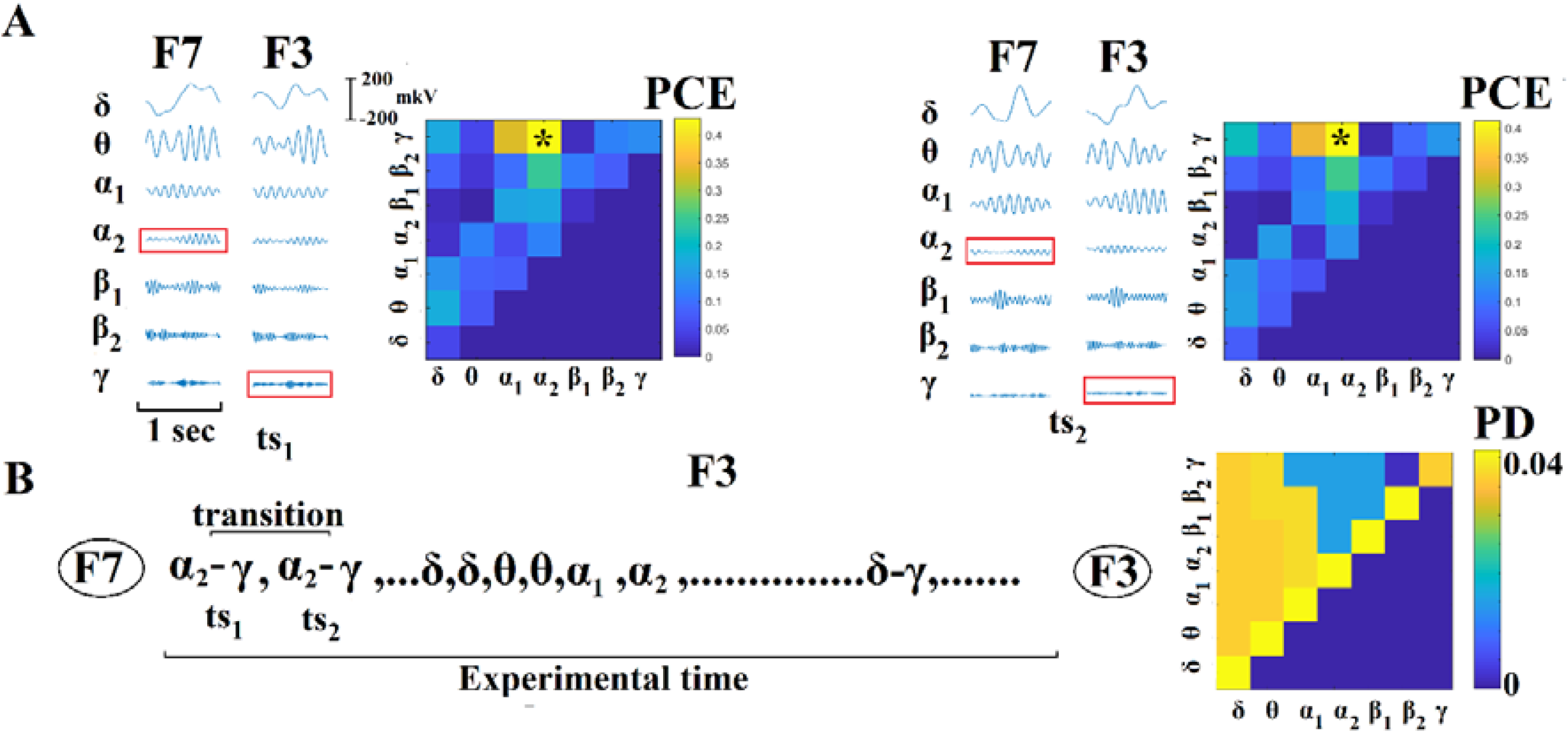
Determining Dominant Intrinsic Coupling Modes (DICM). (A) Schematic illustration of the approach employed to identify the DICM between two EEG sensors (F3 and F7) for two consecutive temporal segment (ts_1_, ts_2_) during the resting□state EEG activity from the first normal subject. In this example the functional interdependence between band□passed signals from the two sensors was indexed by phase coupling estimation (PCE). In this manner PCE was computed between the two EEG sensors either at same□frequency oscillations (e.g., δ to δ) or between different frequencies (e.g., δ to θ). Statistical filtering, using surrogate data approach was employed to assess whether each PCE value was significantly different than chance. During ts_1_ the DICM reflected significant phase locking between α_2_ and γ oscillations (indicated by red rectangles) while during in ts_2_ the dominant interaction was remain stable. (B) Burst of DICM between the two sensors. This packeting can be thought to associate the “letters” contained in the DICM series to form a neural “word.”, a possible integration of many DICM (Buzsaki & Watscon, 2012). For the first pair of ts_1–2_, we illustrated the statibility of DICM for the first pair and a transition between δ - θ and θ-α_1_ is defined for F7□F3 EEG pair important for the estimation of FI (see section 2.3). Based on the DICM time series between every pair of sensors, FI was estimated by counting how many times the DICM changed between consecutive temporal segments divided by the total number of temporal segments −1. FI ranged within [0,1] where higher values are interpreted as higher flexibility. The comodulogram on the right demonstrates the probability distribution (PD) of the DICM across temporal segments for the EEG sensor pair F7-F3. The outcome of this approach is the construction of 2 DFCG or 3D tensors of size [1196 x 16 x 16].

In the present study, we adapted the debiased version of PAC as presented in van Driel et al., (2015).

### 2.3 Semantic Features Derived from the evolution of DICM

This section describes the semantic features that can be defined by analysing the 2^nd^ 3D tensor that preserves the DICM across spatio-temporal space.

#### 2.3.1 Flexibility Index (FI)

Based on the 2nd 3D tensor DIFCG that keeps the information of the DICM per pair of sensors and across time, we estimated the transition rate for every possible pair of sensors. The previously defined estimator was called flexibility index (FI) and quantified how many times every pair of sensors changes DICM across experimental time (Dimitriadis and Salis,2017b; Dimitriadis,2018c). This metric will called hereafter FI^DICM^ which is defined as:

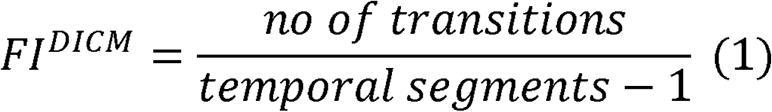

FI^DICM^ gets higher values for higher “jumps” of DICM between a pair of EEG sensors across experimental time. Figure 2b illustrates how FI^DICM^ is estimated for F3-F7 EEG pair. This approach leads to a 16 x 16 matrix per subject or 16^2^ – 16 =140 FI features.

#### 2.3.2 Spatiotemporal distribution of DICM— comodulograms

Based on the 2nd 3D DIFCG that keeps the semantic information of the preferred dominant coupling mode, we can tabulate in a frequencies × frequencies matrix the probability distribution (PD) of observing each of the DICM frequencies across 7 (intra-frequency) + 21 (cross-frequency coupling) = 28 possible coupling modes and exploring their spatio-temporal distribution.

The spatiotemporal PD tabulated in a matrix is called hereafter comodulogram and an example is demonstrated in Fig.2b (Antonakakis et al., 2016,2017a,b; Dimitriadis and Salis,2017b,c; Dimitriadis,2018a,c,d). PD is a new vector of features with dimension 1 x 21 possible DICM.

### 2.4 Low and High-order DFCG

In section 2.2, we described the construction of DIFCG also called as low-order DIFCG (LO-DIFCG). Here, we will define the construction of the high-order DIFCG (HO-DIFCG). The HO-DFCG was constructed by estimating the ‘cosine’ similarity between every pair of the time-series that each one describes the fluctuation of functional strength of a pair of EEG sensors. Here, we quantified the cosine similarity of the fluctuations of PCE by taking into consideration a temporal window of 10 temporal segments sampled from the LO-DIFCG with moving step equals to 1. This means that the 1^st^ temporal dimension of the HO-DFCG is equal to 1187. A LO-FCG has dimensions of 16 x16 while a HO-FCG has dimensions 240 x 240. LO-DIFCG has dimensions of 1196 x 16 x16 while HO-DIFCG has dimensions of 1187 x 240 x 240. Afterward, we topologically filtered both LO and HO-FCG with our OMST method at every temporal segment (Dimitriadis et al., 2017a,c; see supp. Material for further details)^1^.

Practically, HO-DIFCG captures the level of co-fluctuation of functional coupling across EEG-pairs across experimental time. A high cosine similarity value between two pairs of EEG-pairs is interpreted as a high temporal co-fluctuation behaviour.

Fig.3A illustrates the first 20 temporal segments of the LO-DIFCG starting with a full-weighted FCG, a statistical filtering version of FCG (Fig.3B) and its normalized laplacian transformation (see next section) (Fig.3C). Fig.3D illustrates how we constructed in a dynamic way the HO-DIFCG from the LO-DIFCG. Similarly to Fig.3A-C, in Fg.3E, we illustrated the first 20 temporal segments of the HO-DIFCG and its pre-processing steps (Fig.3E-F).

**Figure 3.**
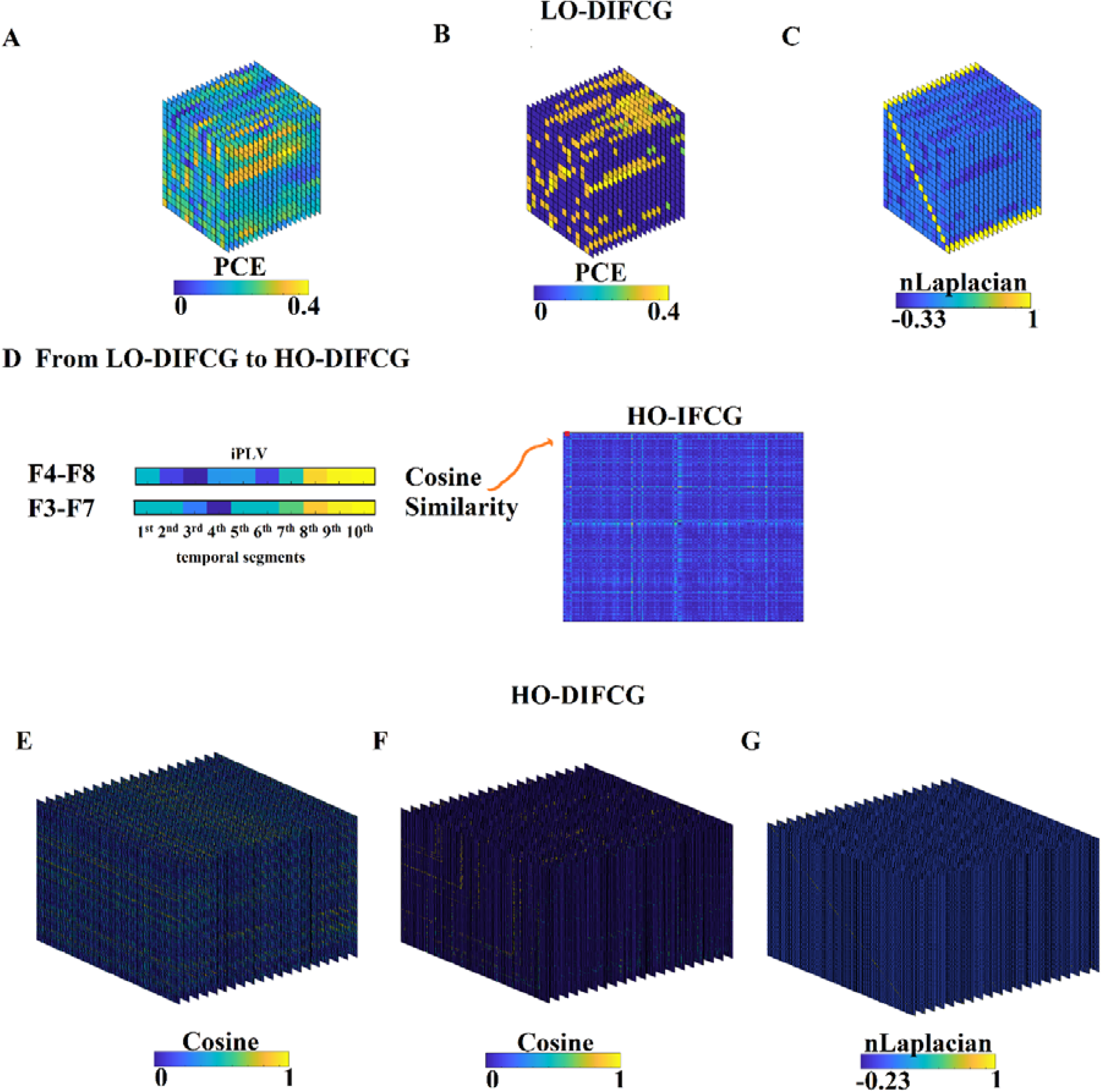
Pre-processing steps of a) LO and b) HO-DIFCG.

#### LO-DIFCG

A. Fully-weighted LO-DIFCG
B. Topological (OMST) and statistically filtered (surrogates) LO-DIFCG
C. NLaplacian transformation of LO-DIFCG demonstrated in (B)

#### From LO-DIFCG to HO-DIFCG

A. We illustrated how HO-DFCG is constructed from LO-DIFCG. We first took the first 10 PCE values from the first 10 temporal-segments from LO-DIFCG (A) for EEG pairs F4-F8 and F3-F7. Then, we estimated the cosine similarity between the two vectors and we finally tabulated this cosine value into a HO-IFCG. The whole procedure was followed for every possible pair of EEG pairs which finally formulated a fully-weighted HO-IFCG. Repeating the same procedure employing width of temporal window that encapsulates 10 temporal segments form the LO-DFCG and re-estimating the cosine similarity from the updated vectors of PCE values, we constructed the HO-DIFCG (E).

#### HO-DIFCG

(E) Fully-weighted HO-DIFCG
(F) Topological (OMST) and statistically filtered (surrogates) HO-DIFCG
(G)NLaplacian transformation of HO-DIFCG demonstrated in (F)

First column demonstrates the full-weighted LO and HO-DIFCG, the second column illustrates the statistically and topologically filtered versions while the third column refers to their normalized laplacian transformations.

### 2.5 Graph Signal Processing of LO and HO-DFCG: Evolving Synchronizability

Given a simple graph FC*G* with *n* vertices, its Laplacian matrix L is defined as (Chung,1997):

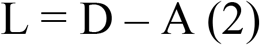

where *D* is the degree matrix and *A* is the adjacency matrix of the graph. Since FCG is a weighted graph, A only contains weights related to the functional strength and its diagonal elements are all 0s (no self functional interactions can be defined for every EEG time series).

In the case of directed graphs, either the in-degree or out-degree might be used, depending on the application. Here, we employed the degree for LO-DIFCG and the in-degree for HO-DIFCG.

The elements of L are given by

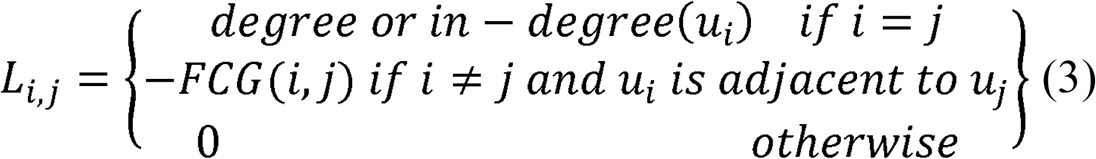

where degree(v_i_) is the degree of the vertex i.

The symmetric normalized Laplacian matrix is defined as:

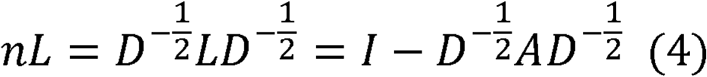

The elements of NL are given by:

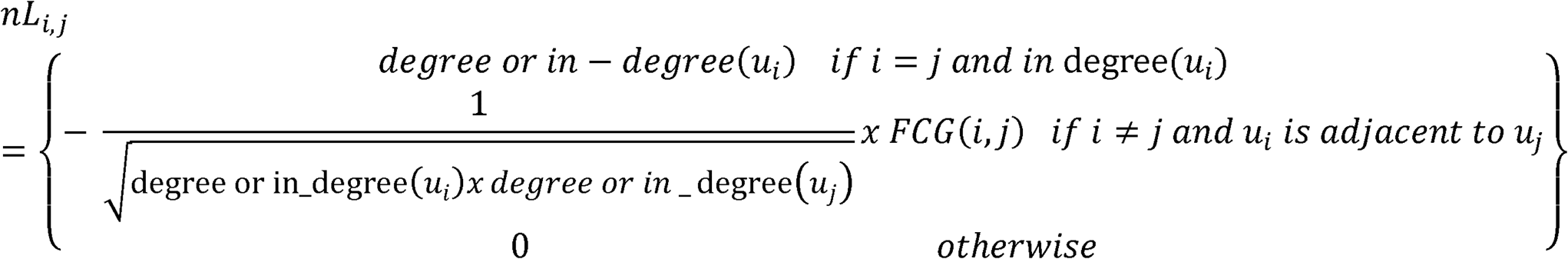

We estimated both L and nL for every snapshot of LO-DIFCG and HO-DIFCG. Then, we calculated the Synchronizability (Syn) index which is defined as the eigenratio of the largest eigenvalue divided by the second smallest one. The analysis has been realized for every quasi-static FCG producing a 2 x 2 network metric time series (NMTS^Syn^) per subject related to {LO and HO-DIFCG} x {L and nL}. Fig.4 demonstrates the four NMTS^Syn^ from the first HC and ScZ subjects.

**Fig. 4.**
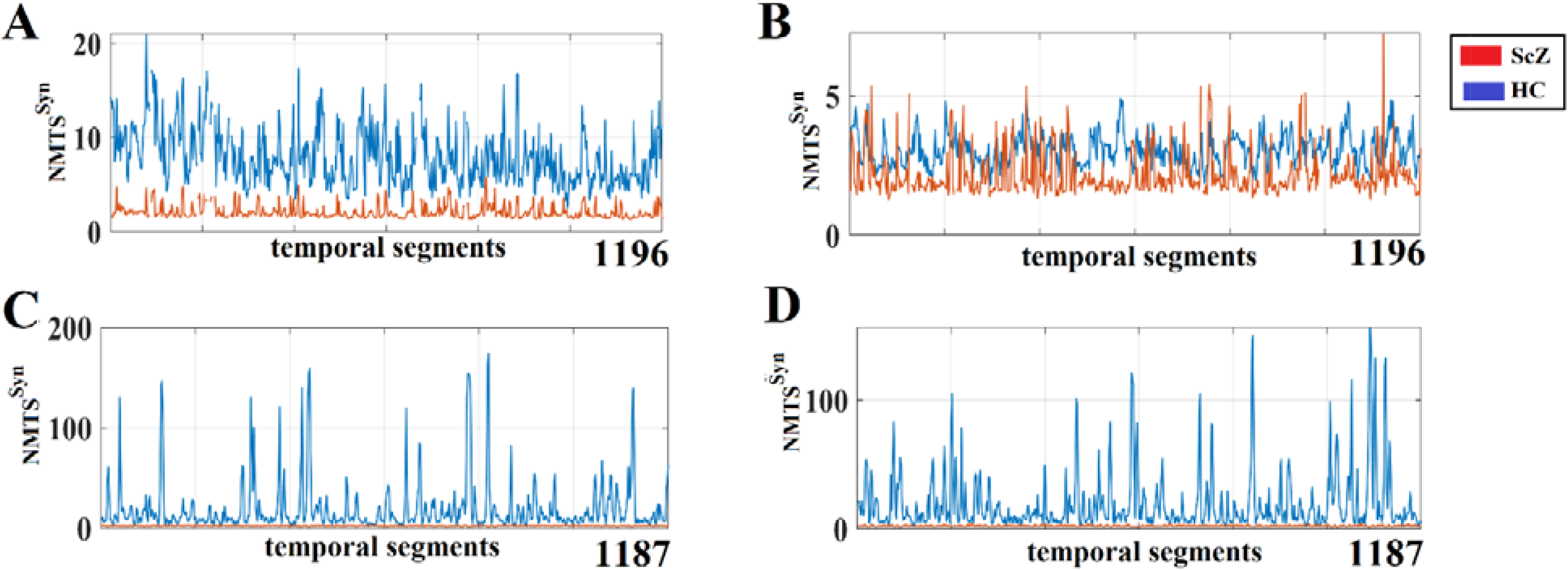
Illustration of representative NMTS^Syn^ from both groups in the four cases. **A)** Laplacian transformation derived from LO-DIFCG **B)** Normalized Laplacian transformation derived from LO-DIFCG **C)** Laplacian transformation derived from HO-DIFCG **D)** Normalized Laplacian transformation derived from HO-DIFCG

### 2.5 Signal Power Analysis

We estimated the relative power spectrum (RSP) of every EEG sensor and frequency band in both groups. Welch’s algorithm was adopted using MATLAB function pwelch leading to the estimation of power spectrum per frequency band first and then their percentage across total power spectrum leading to RSP. Group statistical analysis across sensors and frequency bands have been performed with Wilcoxon Rank Sum test (p < 0.05, Bonferroni corrected, p’< p/(16*7) where 16 refers to EEG sensors and 7 is the number of the studying frequency bands).

### 2.7 Multi-kernel SVM Classification

We divided the whole analysis of corrected classifying ScZ subjects from HC in two parts. The first analysis focused on the analysis of PD of DICM (comodulograms) and FI derived from the DIFCG that keeps the semantic information of the DIFCG, the dominant intrinsic coupling modes. The second analysis focused on the four NMTS^Syn^ derived from LO and HO-DIFCG and eigenanalysis of L and NL.

We hypothesized that PD and FI and the four NMTS^Syn^ provide complementary information to each other. To improve potential classification performance, we decided to fuse those features to further improve the classification performance. Vectorizing the features is not an optimal approach. A kernel-based feature combination using multi-kernel learning offers an alternative approach by weighting differently evert feature derived from a different but complementary procedure (Jie et al., 2015). Such an approach may provide a better way to integrate the features derived from different approaches and types of DIFCG networks.

We adopt a multi-kernel learning to fuse the features by a linear combination of kernels that are estimated from the PD (21 features) and FI (240 features) and by LO and HO-DIFCG, the four NMTS^Syn^. An SVM classifier with a linear kernel *K*(**a**, **b**)□=□**a**^*T*^ **b** based on LIBSVM (Chang and Lin,2011) was employed for the multi-kernel learning based classification based on PD and FI features. The features were first normalized to make sure that all the features are comparable. A linear kernel was calculated across subjects separately for PD and FI. A composite kernel through an optimal linear combination of the multiple kernels was adopted as an optimal fusion of the features.

Particularly for the four NMTS^Syn^, we adopted as a proper kernel tailored to NMTS^Syn^ based on dynamic time wrapping which is a proper distance metric between the NMTS^Syn^ and also Kullback-Leibder divergence histogram distance metric.

We classified every subject with SVM with the composite kernel following a 5-fold cross-validation (CV) scheme 100 runs.

Specifically, for the semantic approach using PD and FI, we adapted as a feature selection algorithm, the infinite feature selection algorithm (Roffo et al., 2015). Feature selection has been applied within every fold of the 5 CV and across runs. From the ranking of 261 features (21: PD + 240: FI) within every fold, we kept the first 15 ranked features. We scored the features that were selected in the first 15^th^ across the 100 runs and 5 folds.

Fig.4 illustrates representative NMTS^Syn^ from the first subject of each group extracted from both LO and HO-DIFCG and also from both L and NL transformations.

## 3. Results

### 3.1 Contribution of NMTS^Syn^ to High Classification Performance of Sch Subjects

Tables 1 summarizes the evaluation of classification performance based on the four NMTS^Syn^, dynamic time wrapping metric and Kullback-Leibder divergence histogram distance metric in the four possible pairs of combinations. The highest accuracy succeeded in LO-DIFCG by combining both L and NL transformations and also by combing the laplacian transformation of LO and HO-DIFCG.

**Table 1.** Evaluation of Classification performance based on NMTS^Syn^, dynamic time wrapping and a multiple-kernel SVM approach.

|  | Accuracy | Sensitivity | Specificity |
| --- | --- | --- | --- |
| **LO-DIFCG (L+NL)** | 0.98 ± 0.0052 | 0.97 ± 0.0052 | 1.00 ± 0.00 |
| **HO-DIFCG (L+NL)** | 0.90 ± 0.02 | 0.95 ± 0.02 | 0.96 ± 0.03 |
| **HO and LO-DIFCG (L)** | 0.98 ± 0.052 | 0.97 ± 0.052 | 0.98 ± 0.01 |
| **HO and LO-DIFCG (NL)** | 0.90 ± 0.02 | 0.95 ± 0.01 | 0.96 ± 0.01 |

Similarly, Tables 2 summarizes the evaluation of classification performance based on the four NMTS^Syn^ and the Kullback-Leibder divergence histogram distance metric in the four possible pairs of combinations. The highest accuracy succeeded in three out of four combinations with the exception of the normalized laplacian transformation of LO and HO-DIFCG (Table 2).

**Table 2.** Evaluation of Classification performance based on NMTS^Syn^, Kullback-Leibder divergence histogram distance metric and a multiple-kernel SVM approach.

|  | Accuracy | Sensitivity | Specificity |
| --- | --- | --- | --- |
| **LO-DIFCG (L+NL)** | 0.97 ± 0.0052 | 0.96 ± 0.01 | 0.97 ± 0.01 |
| **HO-DIFCG (L+NL)** | 0.98 ± 0.01 | 0.97 ± 0.01 | 0.96 ± 0.01 |
| **HO and LO-DIFCG (L)** | 0.98 ± 0.052 | 0.97 ± 0.052 | 0.98 ± 0.01 |
| **HO and LO-DIFCG (NL)** | 0.87 ± 0.03 | 0.90 ± 0.02 | 0.92 ± 0.02 |

### 3.2 Contribution of Semantic Features based on DICM model for High Classification Performance of Sch Subjects

Our machine learning strategy revealed 25 consistent features among FI and PD of DICM. Fig.5 (a-d) illustrates the spatial distribution of the FI across every possible pair of EEG sensors for HC and ScZ group, correspondingly. Feature selection strategy revealed 10 FI features that are located over fronto–temporo–parietal network and denoted with ‘*’. Figure 6 (e-f) illustrates the group□averaged comodulograms for HC and ScZ group, correspondingly. 15 more features were selected that referred to the PD across space and time of θ-θ α_1_-α_1_ α_2_-α_2_ β_2_-β_2_ (intra-frequency coupling modes) and the following cross-frequency coupling modes: δ-α_2_ δ-β_1_ δ-β_2_ (δ modulating frequency), θ-β_1_ θ-β_2_ (θ modulating frequency), α_1_-β_1_ α_1_-β_2_ α_1_-γ (α_1_ modulating frequency), α_2_-β_1_ α_2_-β_2_ α_2_-γ (α_2_ modulating frequency)intrinsic coupling modes. ScZ patients demonstrated lower PD values compared to HC. The classification performance based on selected FI+PD features was absolute (100%). Table 3 tabulates the performance for each set of features and also their integration.

**Fig. 5.**
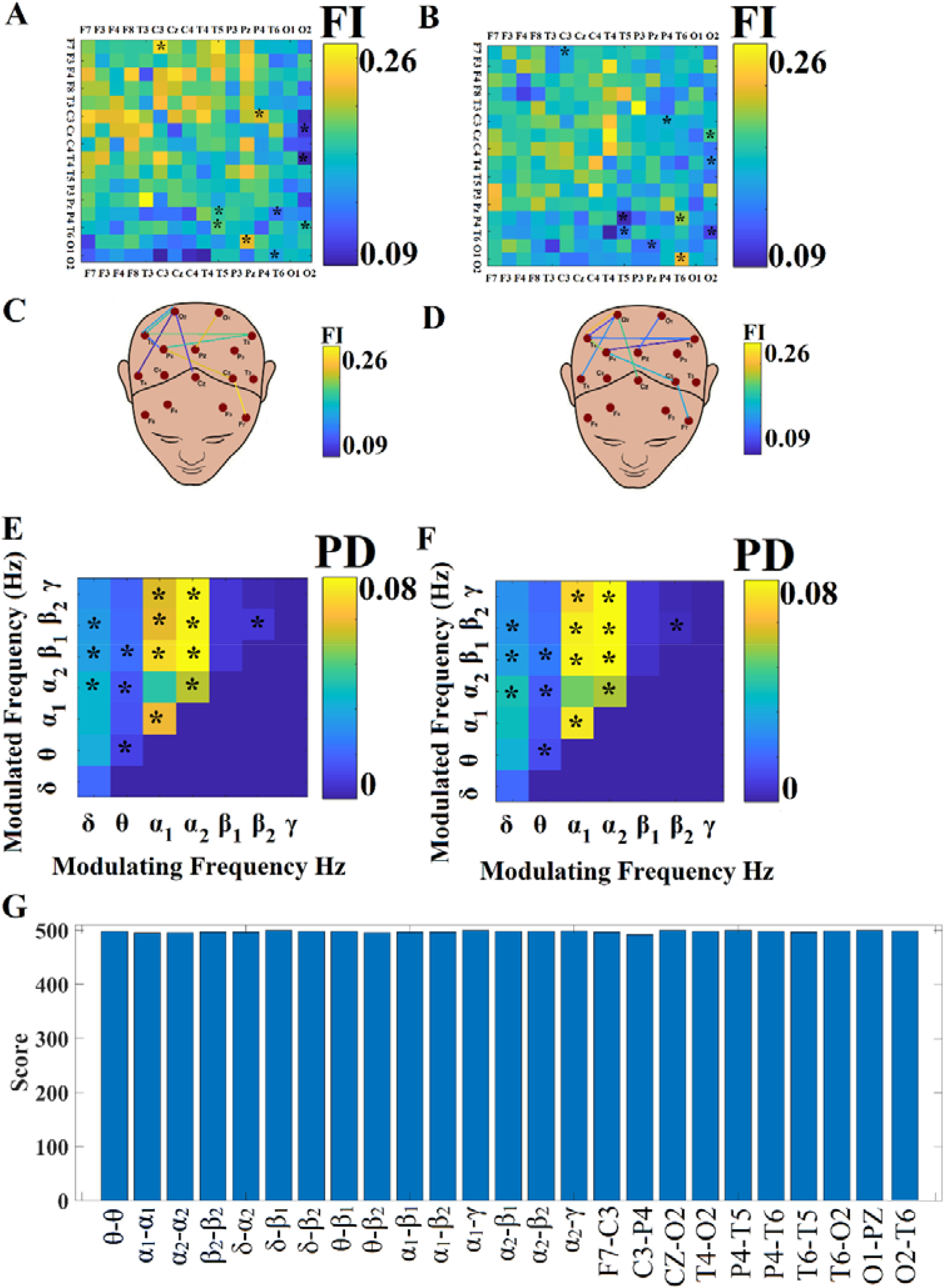
Group□Averaged Flexibility Index (FI) and Comodulograms that tabulate the PD of spatio-temporally DICM. (A-B) Group□averaged FI for healthy control (HC) (A) and schizophrenic (ScH) group (B). (C-D) Group□averaged comodulograms for HC (C) and ScZ group (E-F) probability distribution of DICM across space and time averaged across subjects in (E) HC group and in (F) ScZ group. (G) Total score of the selected features demonstrated in a – f across 100 runs x 5-fold CV = 500 times. Every FI and PD has been selected via the feature selection algorithm is denoted with ‘*’.

**Table 3.** Evaluation of Classification performance based the semantic features pool of PD and FI and on a multiple-kernel SVM approach.

|  | Accuracy | Sensitivity | Specificity |
| --- | --- | --- | --- |
| **PD** | 1.00 0.00 | 1.00 0.00 | 1.00 0.00 |
| **FI** | 1.00 0.00 | 1.00 0.00 | 1.00 0.00 |
| **PD+FI** | 1.00 0.00 | 1.00 0.00 | 1.00 0.00 |

**Figure 6.**
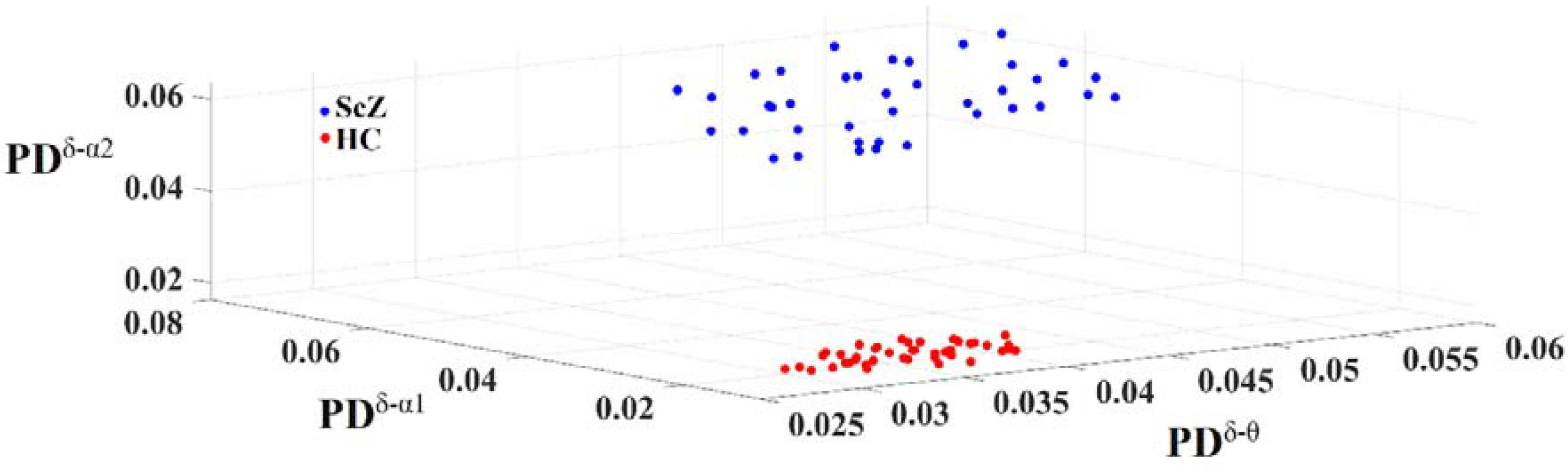
Discriminative power of PD for δ□θ, δ□α_1_, δ□α_2_ cross-frequency pairs. Every dot corresponds to every single subject.

Finally, Fig.5(f) demonstrates the Score of the selected features across 100 runs x 5 – fold CV which refers to the number of times a feature is selected as informative to discriminate HC from ScZ group.

Fig.6 illustrates the high discriminative power of PD for δ□θ, δ□α_1_, δ□α_2_, in a 3D plot. The two groups allocate a different sub-space which further supported our absolute accuracy.

### 3.3 Relative Signal Power in HC and ScZ subjects

Statistical analysis of RSP between the two groups across EEG sensor space and frequency bands revealed significant higher δ RSP for ScZ compared to HC in O1 and O2 EEG sensors. Fig.7 illustrates a global tendency (no statistical significant) of higher RSP values in δ and θ frequencies in ScZ compared to HC subjects.

**Figure 7.**
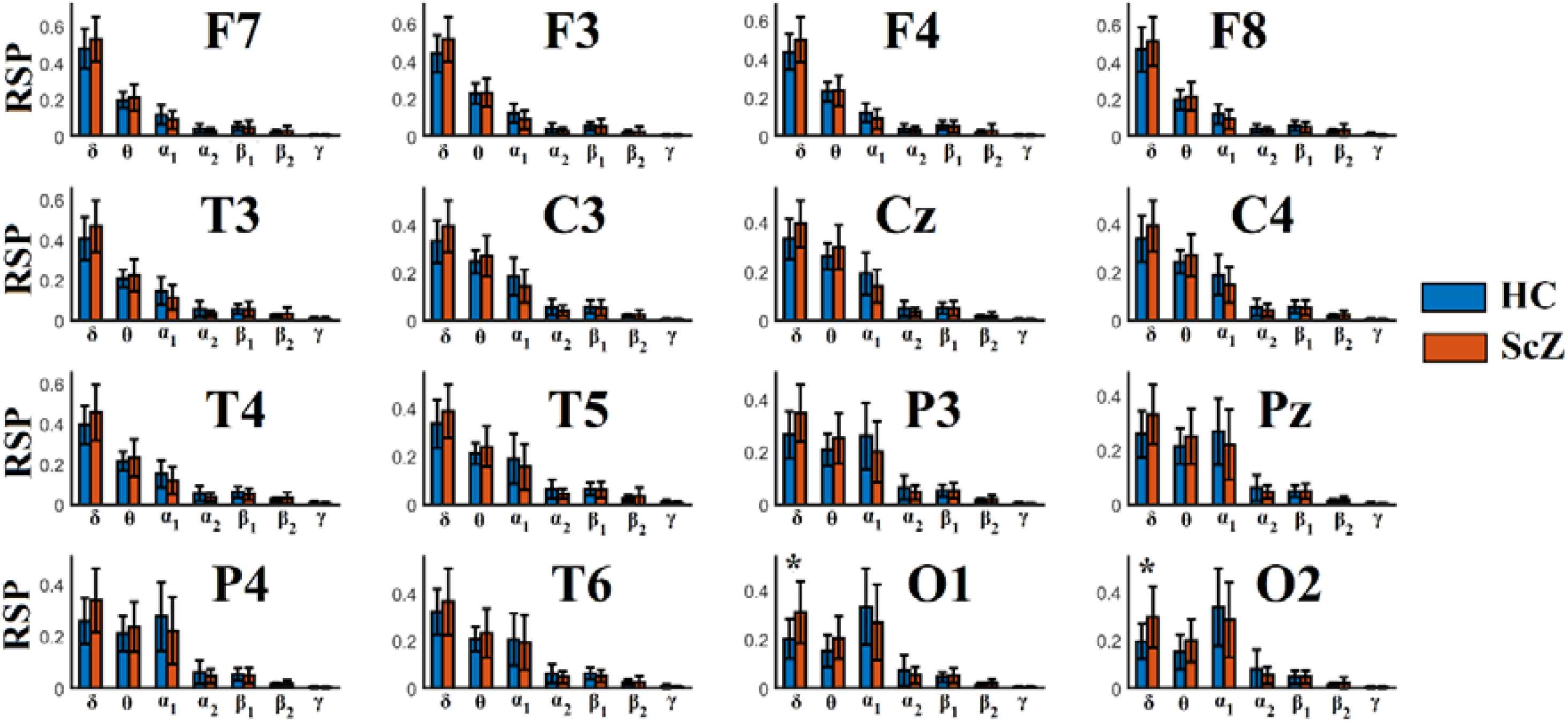
(Relative Signal Power (RSP) across EEG sensor space and frequency bands in
HC and ScZ patients. Our analysis revealed significant higher δ RSP for ScZ compared to HC in O1 and O2 EEG sensors. (* Wilcoxon Rank Sum test (p < 0.01, Bonferroni corrected, p’< p/(16*7) where 16 refers to EEG sensors and 7 to the number of the studying frequency bands).

## 4. Discussion

In the present study, we investigated the contribution of graph signal processing over LO and HO-DIFCG and semantic information derived from our DICM model to the diagnosis of schizophrenia. Following a data-driven model from EEG recordings up to the set of novel chronnectomics metrics, we succeeded to discriminate HC from ScZ patients with absolute accuracy. Laplacian transformations of LO and HO-DIFCG gives the baton to the estimation of the Synchronizability giving rise to four representative NMTS^Syn^ per subject. DICM model extracts 261 features explicitly refer to PD (21 features) and FI (240 features) per subject. Following an integrated multi-kernel learning SVM approach, we succeeded to discriminate ScZ patients from HC with 98% using NMTS^Syn^ and 100% employing PD and FI.

It is important to draw attention to neuroscientists working on functional neuroimaging the significance of exploring simultaneously the spatio-temporal behaviour of functional strength complementary to the PD of DICM across space and time. Here, we revealed a global temporal hypo-synchronization of ScZ patients compared to HC subjects (Fig.4). This hypo-synchronization in HO-DIFCG can be interpreted as a loss of co-fluctuation property of dynamic functional coupling between pairs of EEG pairs.

Our data-driven methodology that quantified the dynamic synchronizability of both LO and HO-DIFCG revealed whole-brain breakdown for ScZ patients. The classification performance based on NMTS^Syn^ was high reaching 98% in many cases including the combination of NMTS^Syn^ derived from both LO and HO-DIFCG. In the fusion approach (LO – HO-DIFCG), laplacian transformation succeeded better performance than normalized laplacian. However, HO-DIFCG didn’t improve further the very high classification performance in the multiple-kernel approach. It seems that in a few subjects, NMTS^Syn^ didn’t provide complementary information between LO and HO-DIFCG.

Complementary, our analysis revealed a less flexible brain for ScZ patients which is interpreted as less switching functional interactions between dominant intrinsic coupling modes. The spatial distribution of the FI features is extended between EEG sensors located over fronto–temporo–parietal brain areas. Feature selection strategy revealed lower PD values for ScZ patients for eight intrinsic coupling modes: δ□α_2_, δ□β_1_, δ□β_2_, α_1_-β_2_ α_1_-γ,α_2_-β_1_ α_2_-β_2_ α_2_-γ (Fig.5). It is the very first time that a study reported these abnormalities in schizophrenia research under the framework of a unified model of identifying dominant intrinsic coupling modes. These quantitative and qualitative novel features succeeded to absolute discriminate (100%) healthy controls from schizophrenia patients, a result of significant importance for schizophrenia research.

Schizophrenia includes an abnormal functionality of many brain areas including the hippocampus (Tamminga et al.,2010). Hippocampal changes explained memory abnormalities observed in schizophrenia and may affect also other brain areas. Specifically, in ScZ patients, researchers observed an hyper-activity of CA1 (Lodge and Grace,2011). *N*-methyl-D-aspartate receptor (NMDAR) antagonist mimicked many symptoms of schizophrenia (Zhang et al., 2012). Of great importance is the enhancement of δ activity at awake state in schizophrenia, an aberrant functionality that can be explained by NMDAR antagonist action in the thalamus that can stimulate thalamic activity and produce δ oscillations (Sharp and Hendren 2007). Here, we detected significant higher RSP in δ frequency in ScZ patients compared to HC in O1 and O2 EEG sensors (Fig.6).

Many previous studies explored the contribution of both low and high frequency oscillations to explain a range of cognitive deficits in schizophrenia modulated with specific frequency content (Moran and Hong,2011). A large study explored the hierarchical organization of γ and θ oscillations in schizophrenia during a 40 Hz auditory steady-state stimulation. They found out that ScZ patients revealed a reduced γ inter-trial phase coherence, increased θ amplitude, and normal θ-phase γ-amplitude cross-frequency coupling (Kirihara et al., 2012). In the present study and under the framework of DICM model, we didn’t detect also significant differences of the PD of θ-phase γ-amplitude cross-frequency coupling between the two groups.

It is important to state here that this is the very first study that explored simultaneously both intra-frequency and cross-frequency interactions in Schizophrenia under DICM model. Under this framework, we can decipher the multiplexity of complex electrophysiological aberrant connectivity observed in schizophrenic patients by integrating the available intrinsic coupling modes.

Our DICM model untangled the multiplex syntactic rules which are important for the exchange of information across the cortex (Buzsaki & Watscon, 2012). DICM model hypothesizes that the dynamic reconfiguration of dominant intrinsic coupling modes can capture the multiplexity and complexity of functional brain connectivity during both spontaneous and cognitive tasks. Additionally, the flexibility of this reconfiguration can be linked to the multiplexity of brain functionality and can be estimated via FI. FI can be seen as a reflex index that describes the readiness of the brain to respond to new stimuli.

It would be interesting to follow the same methodology during cognitive tasks and also on the source level in order to get the advantage of animal models and fMRI studies that revealed the mechanistic explanation of aberrant networks in schizophrenia (Hunt et al., 2017).

## 5. Limitations

The whole study is unique, innovative and pioneering in terms of analytic pathway and scientific results. However, there are two basic limitations. The first refers to the interpretation of the results on the EEG surface level instead of virtual cortical sources. It would be interesting to follow the same methodological approach in an EEG study recorded healthy controls and schizophrenic patients using a large number of EEG net sensors that support the source reconstruction approach. The second drawback is the limitation of the adopted free database to EEG recordings only without any access to neuropsychological assessment battery. This missing part prevented us to correlate the novel chronnectomic and semantic features with trivial neuropsychological estimates.

## Acknowledgments

SID was supported by MRC grant MR/K004360/1 (Behavioral and Neurophysiological Effects of Schizophrenia Risk Genes: A Multi-locus, Pathway Based Approach). SID is also supported by a MARIE-CURIE COFUND EU-UK Research Fellowship. We would like to acknowledge RCUK of Cardiff University and Wellcome Trust for covering the publication fee.

This manuscript has been released as a Pre-Print at: https://www.biorxiv.org/content/10.1101/551671v1 with **doi:** https://doi.org/10.1101/551671

## Footnotes

1 https://github.com/stdimitr/topological_filtering_networks

